# Phase Separation in Mixtures of Prion-Like Low Complexity Domains is Driven by the Interplay of Homotypic and Heterotypic Interactions

**DOI:** 10.1101/2023.03.15.532828

**Authors:** Mina Farag, Wade M. Borcherds, Anne Bremer, Tanja Mittag, Rohit V. Pappu

## Abstract

Prion-like low-complexity domains (PLCDs) are involved in the formation and regulation of distinct biomolecular condensates that form via coupled associative and segregative phase transitions. We previously deciphered how evolutionarily conserved sequence features drive phase separation of PLCDs through homotypic interactions. However, condensates typically encompass a diverse mixture of proteins with PLCDs. Here, we combine simulations and experiments to study mixtures of PLCDs from two RNA binding proteins namely, hnRNPA1 and FUS. We find that 1:1 mixtures of the A1-LCD and FUS-LCD undergo phase separation more readily than either of the PLCDs on their own. The enhanced driving forces for phase separation of mixtures of A1-LCD and FUS-LCD arise partly from complementary electrostatic interactions between the two proteins. This complex coacervation-like mechanism adds to complementary interactions among aromatic residues. Further, tie line analysis shows that stoichiometric ratios of different components and their sequence-encoded interactions jointly contribute to the driving forces for condensate formation. These results highlight how expression levels might be tuned to regulate the driving forces for condensate formation *in vivo*. Simulations also show that the organization of PLCDs within condensates deviates from expectations based on random mixture models. Instead, spatial organization within condensates will reflect the relative strengths of homotypic versus heterotypic interactions. We also uncover rules for how interaction strengths and sequence lengths modulate conformational preferences of molecules at interfaces of condensates formed by mixtures of proteins. Overall, our findings emphasize the network-like organization of molecules within multicomponent condensates, and the distinctive, composition-specific conformational features of condensate interfaces.

**Significance Statement:** Biomolecular condensates are mixtures of different protein and nucleic acid molecules that organize biochemical reactions in cells. Much of what we know about how condensates form comes from studies of phase transitions of individual components of condensates. Here, we report results from studies of phase transitions of mixtures of archetypal protein domains that feature in distinct condensates. Our investigations, aided by a blend of computations and experiments, show that the phase transitions of mixtures are governed by a complex interplay of homotypic and heterotypic interactions. The results point to how expression levels of different protein components can be tuned in cells to modulate internal structures, compositions, and interfaces of condensates, thus affording distinct ways to control the functions of condensates.

---

Biomolecular condensates are membraneless bodies that provide spatial and temporal control over cell signaling and cellular responses to various stresses (1–7). Functional roles for condensates have been implicated in transcriptional regulation (8–14), cytosolic and nuclear stress responses (3, 15–21), trafficking of cellular components (22, 23), RNA regulation and processing (24–28), mechanotransduction (29–31), and protein quality control (32–36). The working hypothesis, based on a growing corpus of data, is that condensates form via spontaneous and driven phase transitions of networks of multivalent biomacromolecules (1, 37, 38). The relevant processes involve a coupling of associative and segregative phase transitions (39). These include processes such as phase separation coupled to percolation (PSCP) (39–41) and complex coacervation (42–44). Multivalent proteins that scaffold and drive phase transitions encompass different numbers and types of oligomerization and substrate binding domains (41). Most, although not all protein scaffolds also feature intrinsically disordered regions (IDRs) that drive or modulate phase separation of protein scaffolds through a blend of homotypic and heterotypic interactions (45). Here, homotypic, and heterotypic interactions refer to intermolecular interactions between the same versus different molecules, respectively. This concept can be extended to distinguish interactions between the same versus different motifs on molecules (46).

IDRs that drive condensate formation include a family of sequences referred to as prionlike low-complexity domains (PLCDs) (47, 48). These domains are readily recognizable based on their overall compositional biases (49). Our analysis of 89 condensate-associated PLCDs drawn from the human proteome shows that on average, 50-60% of the amino acid residues within PLCDs are polar (Gly, Ser, Thr, Gln, and Asn), ~10% are aromatic (Tyr, Phe, Trp, and His), ~13% are Pro, fewer than 5% are charged, and the remaining residues are aliphatic (**Fig. 1A**).

**Fig. 1:**
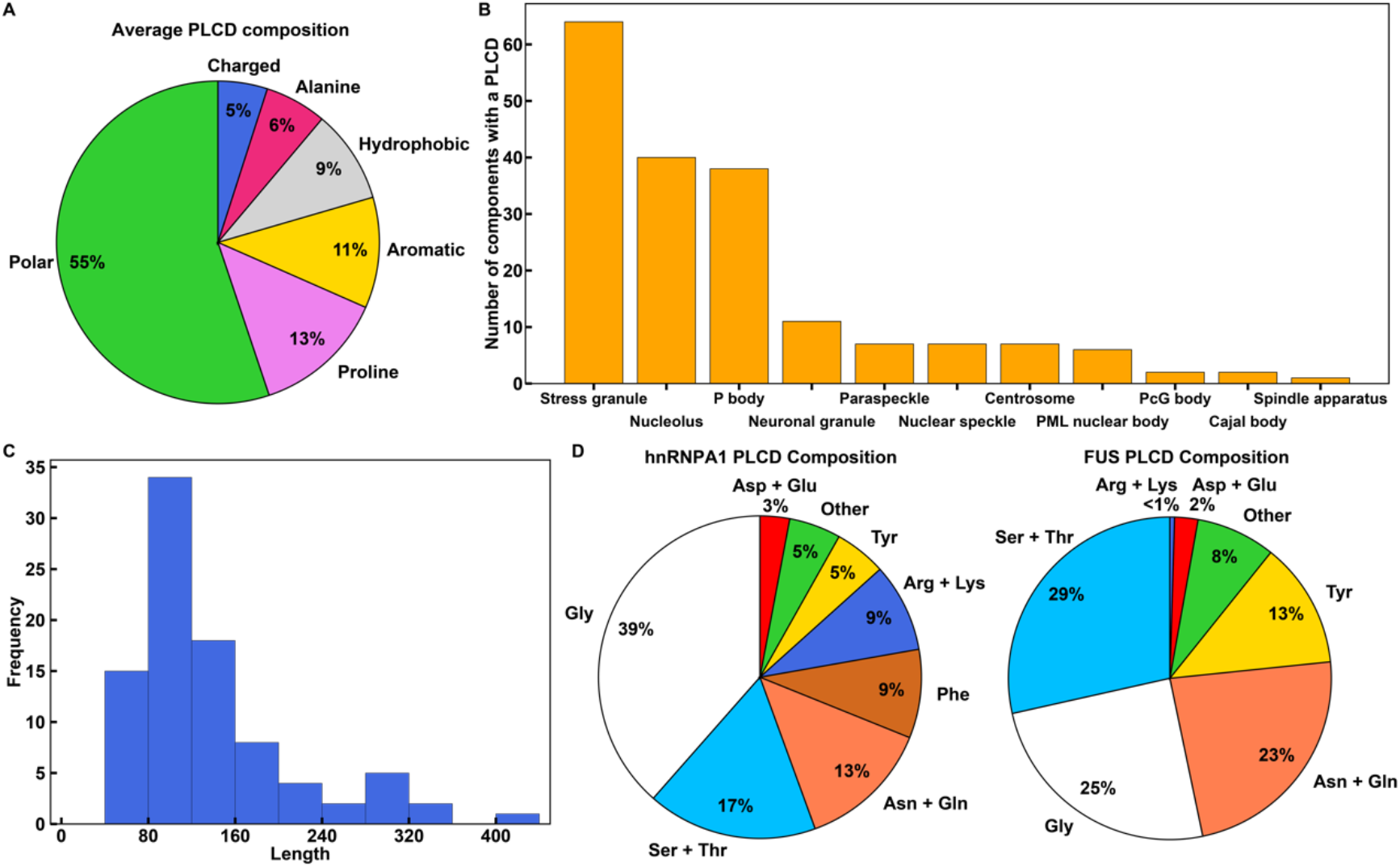
PLCDs of similar albeit non-identical compositions are prevalent in distinct biomolecular condensates. **(A)** Average compositional profile of 89 PLCDs found within cellular condensates in *Homo sapiens.* Here, “Polar” includes Gly, Ser, Thr, Asn, Gln, and Cys. “Aromatic” includes Phe, Tyr, Trp, and His. “Hydrophobic” includes Leu, Iso, Val, and Met. “Charged” includes Asp, Glu, Arg, and Lys. **(B)** The Distribution of PLCDs across cellular condensates in *Homo sapiens.* The condensates examined have at least 40 known protein components. **(C)** Length distribution of all 89 distinct PLCDs in **(A), (B)**. **(D)** Compositional profiles of PLCDs within human versions of hnRNPA1 and FUS. For the hnRNPA1 PLCD, the “Other” category includes 2.2% Ala, 1.5% Met, and 1.5% Pro. For the FUS PLCD, the “Other” category includes 1.9% Ala, 1.0% Met, and 5.1% Pro.

Recent studies have focused on quantitative assessments of how distinct sequence features contribute to the driving forces for phase transitions of PLCDs. The stickers-and-spacers model for PSCP (41, 50–52) has provided an organizing framework for uncovering sequence-specific driving forces of PLCDs (53–55). The emerging picture may be summarized as follows: Aromatic residues function as stickers or cohesive motifs (53, 54). Physical crosslinks among stickers contribute to the networking of PLCDs within condensates (55). The number of aromatic stickers appears to be one of the main determinants of threshold concentration for phase separation (53, 54). Further, across evolution, stickers are non-randomly and uniformly distributed along linear sequences of PLCDs (53, 56). Polar residues function as spacers interspersed between stickers. However, spacers are not inert entities (54). Instead, their solvation or excluded volumes dictate the extent of coupling between networking, via reversible sticker-sticker crosslinks, and the driving forces for phase separation, via the influence on overall protein solubility (57). For example, increasing the net charge of spacers can have a destabilizing effect on phase separation of PLCDs. Spacers also influence the cooperativity of sticker-sticker interactions (54).

Two recent studies generalized the binary stickers-and-spacers model, showing that Tyr is a stronger sticker than Phe within PLCDs (54, 55). Within PLCDs, Arg plays the role of an auxiliary rather than a primary sticker. This stands in contrast to findings for full-length FET family proteins such as Fused in Sarcoma (FUS), where Tyr and Arg residues are the primary stickers in proteins that encompass both a PLCD and disordered RNA binding domains (52, 58). Accordingly, instead of being immutable (59), the relative strengths of sticker-sticker interactions and the identities of primary versus auxiliary stickers are highly context-specific (39, 60). Context dependence and compositional specificity extends to spacers as well (52). For example, Gly and Ser are non-equivalent as spacers in PLCDs (54, 55). For a given composition of stickers, Gly residues act as spacers that enhance the driving forces for phase separation when compared to Ser and other polar amino acids (54, 55). However, long poly-Gly tracts are generally avoided within PLCDs because these tracts become alternative stickers that can drive fibril formation (61). Evolutionarily, one observes strong correlations between sticker versus spacer identities in PLCDs. For example, the fractions of Tyr and Phe residues as well as Gly and Ser residues tend to be inversely correlated with one another. Additionally, positive correlations were observed between Tyr and Gly contents as well as Phe and Ser contents (54).

The driving forces for phase transitions of individual PLCDs depend on sequence-specific compositional biases (54, 55). Indeed, evolutionarily observed compositional variations can give rise to differences in driving forces for PSCP that can span several orders of magnitude (54). Condensates such as stress granules and P bodies, and nuclear bodies such as nucleoli house ~40 or more known proteins with distinct PLCDs (**Fig. 1B**). Within a condensate, the PLCDs can be quite different from one another. The sequence lengths of PLCDs can vary by a factor of four, with most PLCDs being ~150 residues long (**Fig. 1C**). Therefore, the driving forces for condensate formation in mixtures of PLCDs are of direct relevance for understanding how different PLCDs work together or in opposition to influence condensate formation, internal organization, and interfacial properties.

Here, we report results from studies of condensate formation in mixtures of PLCDs from two proteins, hnRNPA1 (62) and FUS (63). These two proteins are key components of stress granules, paraspeckles, and other condensates (64). Hereafter, we refer to the two PLCDs as A1-LCD and FUS-LCD, respectively. Mutations within both PLCDs are associated with the formation of aberrant stress granules in the context of Amyotrophic Lateral Sclerosis (ALS) (20, 62, 65). The compositional differences between human versions of the two PLCDs are summarized in **Fig. 1D**. These differences translate to a positive net charge per residue (NCPR) for A1-LCD and a negative NCPR for FUS-LCD. In addition to being compositionally different, the two sequences have different lengths, with FUS-LCD being ~1.6 times longer than A1-LCD. However, despite being shorter than FUS-LCD, the driving forces for PSCP, as measured by the temperature- and solutioncondition-dependent saturation concentrations (*c*_sat_), are stronger for A1-LCD when compared to FUS-LCD (55). Given these differences, we focused on understanding how the interplay of homotypic and heterotypic interactions influences the driving forces for condensate formation in mixtures of A1-LCD and FUS-LCD molecules. For this, we deployed a recently developed coarsegrained model to simulate temperature-dependent phase transitions of mixtures of A1-LCD and FUS-LCD (55). In the model, each PLCD residue is modeled as a single bead with distinct interbead interactions. The simulations were performed using LaSSI (51), which is a lattice-based Monte Carlo simulation engine.

We find that heterotypic interactions are the dominant contributors to condensate formation in 1:1 mixtures of A1-LCD and FUS-LCD. This leads to two-component phase diagrams with distinctive shapes and slopes for tie lines. We demonstrate the accuracy of our computational results using *in vitro* experiments that leverage a novel analytical HPLC-based method for measuring phase diagrams in multi-component mixtures (66). Further, through additional simulations, we uncover general rules for how the interplay between homotypic and heterotypic interactions influences the internal organization and interfacial properties of multicomponent condensates.

## Results

### Heterotypic interactions enhance the driving forces for phase separation of mixtures of A1-LCD and FUS-LCD

We performed a series of simulations for mixtures of A1-LCD and FUS-LCD. The sequences of A1-LCD and FUS-LCD are shown in **Fig. 2A**. The total protein concentrations were fixed in the simulations, and the ratios of FUS-LCD-to-A1-LCD were varied from one set of simulations to another. Treating the mixture as a system with one type of macromolecule, we computed binodals in the plane of total protein concentration along the abscissa and temperature along the ordinate. This approach to depicting phase boundaries for mixtures parallels that of Elbaum-Garfinkle et al., (67) and Wei et al., (68). Based on the computed binodals (**Fig. 2B**) we predict that a 1:1 mixture of FUS-LCD and A1-LCD undergoes phase separation at a lower total protein concentration than either FUS-LCD or A1-LCD on its own. This is suggestive of the presence of heterotypic interactions that enhance phase separation.

**Fig. 2:**
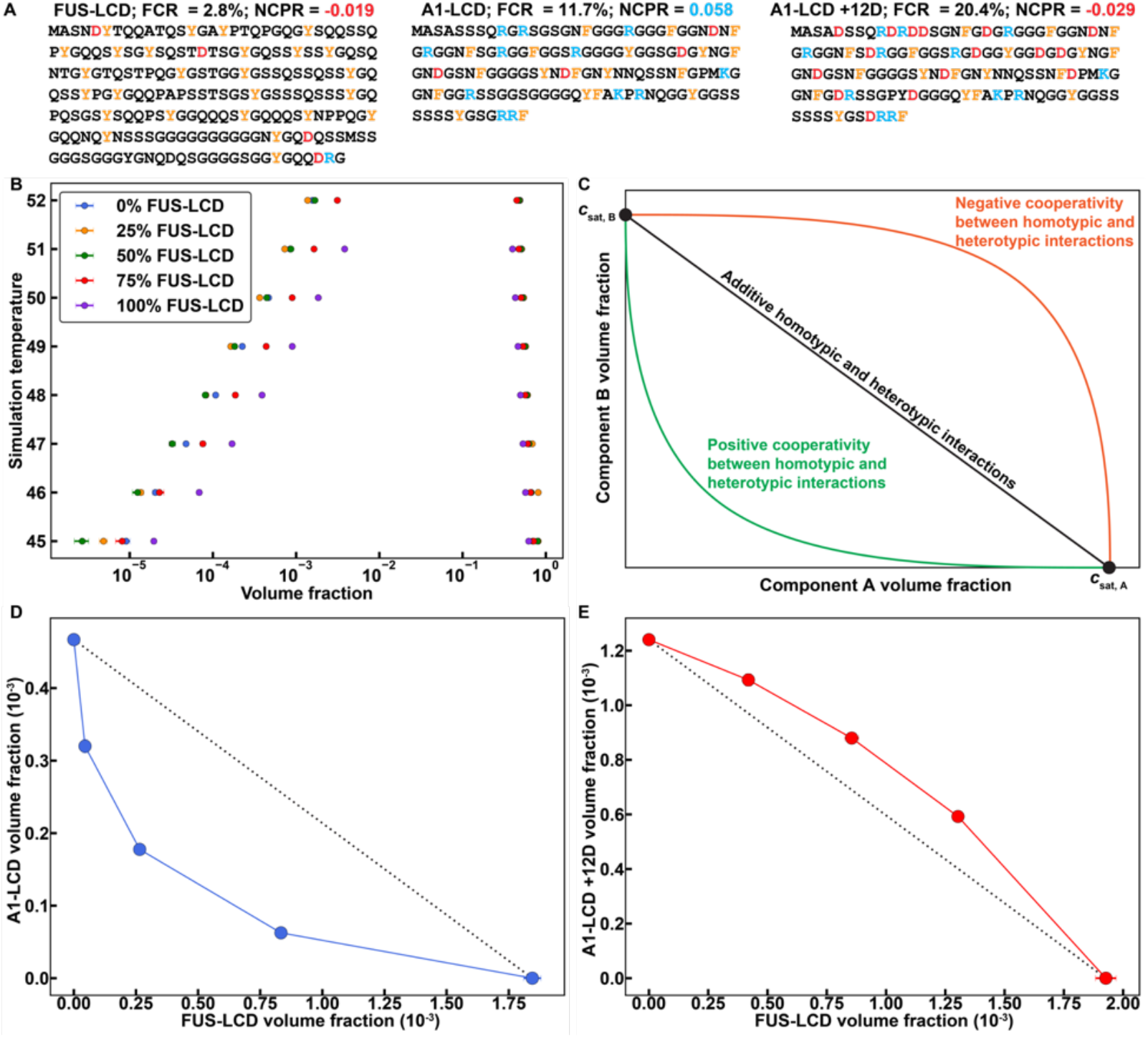
LaSSI simulations predict that phase behaviors of mixtures of PLCDs are governed by the interplay of homotypic and heterotypic interactions. **(A)** Details of the amino acid sequences of FUS-LCD, A1-LCD, and A1-LCD +12D, including the fraction of charged residues (FCR) and the net charge per residue (NCPR). FUS-LCD has 214 residues, whereas A1-LCD and A1-LCD +12D each have 135 residues. **(B)** Computed binodals of mixtures of FUS-LCD and A1-LCD. Volume fractions correspond to the sum of the FUS-LCD and A1-LCD volume fractions. Percentages described in the legend indicate the relative ratio of FUS-LCD to A1-LCD. **(C)** Schematic depicting the expected shape of the binodal depending on the interplay between homotypic vs. heterotypic interactions. **(D),(E)** Computed 2-component binodals of mixtures of FUS-LCD and **(D)** A1-LCD at a fixed simulation temperature of 49 or **(E)** A1-LCD +12D at a fixed simulation temperature of 50. Black dashed lines connect the intrinsic *c*_sat_ of FUS-LCD to the intrinsic *c*_sat_ of A1-LCD or A1-LCD +12D and are shown to indicate the expected binodal shape if heterotypic interactions are on par with homotypic interactions. Error bars in **(B)**, **(D)**, and **(E)** are standard errors about the mean across 5 replicates. Where error bars are invisible, they are smaller than the marker size.

To better understand the interplay between homotypic and heterotypic interactions, we recast the results as a two-dimensional phase diagram at a fixed temperature, where each axis is defined by the concentration of one protein. In **Fig. 2C** we show expectations for the dilute arms of two-dimensional phase diagrams for a system of two components that undergo co-phase separation from solution.

At the temperature of interest, we shall denote the saturation concentration of A1-LCD in the absence of FUS-LCD as *c*_sat,A1_. Likewise, at the same temperature, the saturation concentration of FUS-LCD in the absence of A1-LCD is denoted as *c*_sat,FUS_. If the dilute arm of the twodimensional phase diagram is a *straight line* joining the individual *c*_sat_ values, then the contributions to phase separation of the mixture of PLCDs are purely *additive.* In this case, homotypic and heterotypic interactions make equivalent contributions to the driving forces phase separation and there is no cooperativity in the system. As a result, the total protein concentration in the dilute phase of the two-phase system can be written as: *c*_dilute_ = *c*_sat,A1_ + (1 – *α*)*c*_sat,FUS_, where *a* is the fraction of A1-LCD molecules in the system; when *a* = 1, *c*_dilute_ = *c*_sat,A1_, and when *a* = 0, *c*_dilute_ = *c*_sat,FUS_. If the computed or measured dilute arm of the two-dimensional phase diagram is *concave*, then heterotypic interactions *enhance* the driving forces for phase separation. In this scenario, there is *positive cooperativity*, whereby phase separation is enhanced by mixing the two components. Conversely, if the dilute arm of the two-dimensional phase diagram is *convex*, then heterotypic interactions *weaken* the driving forces for phase separation. This is a manifestation of *negative cooperativity*, whereby phase separation is weakened in a mixture of the two components. The degree to which the driving forces for phase separation are enhanced or weakened depends on the composition of the mixture i.e., the stoichiometric ratio of the two components.

The results shown in **Fig. 2B** were recast by fixing the simulation temperature and plotting the computed dilute phase concentrations of A1-LCD and FUS-LCD for different stoichiometric ratios (**Fig. 2D**). We observe a concave shape, indicating an enhancement of phase separation via heterotypic interactions. FUS-LCD is negatively charged, and A1-LCD is positively charged. Accordingly, we reasoned that complementary electrostatic interactions are likely to enhance the driving forces for co-phase separation. These interactions are likely to contribute in addition to heterotypic aromatic sticker interactions. To test this hypothesis, we performed simulations for mixtures of FUS-LCD and a variant of A1-LCD denoted as A1-LCD +12D. In this variant, twelve Asp residues were substituted across the sequence, replacing extant spacers, giving rise to a sequence with a net negative charge. Although the aromatic sticker interactions remain unchanged, the electrostatic interactions should be weakened. We reasoned that this would generate a dilute arm with a more convex shape, and this is precisely what we observe (**Fig. 2E**).

### Results from in vitro measurements are in accord with computational predictions

Next, we measured co-phase separation in aqueous mixtures of A1-LCD and FUS-LCD molecules (see Materials and Methods, *SI Appendix* and **Fig. S1**). Diffraction-limited fluorescence microscopy shows that the PLCDs co-localize into the same condensates upon phase separation (**Fig. 3A**). This is true for all concentration ratios studied. We then used our recently described analytical high-performance liquid chromatography (HPLC) method (66) to determine dilute and dense phase concentrations of both species in mixtures with different mass concentration ratios of the two PLCDs.

**Fig. 3:**
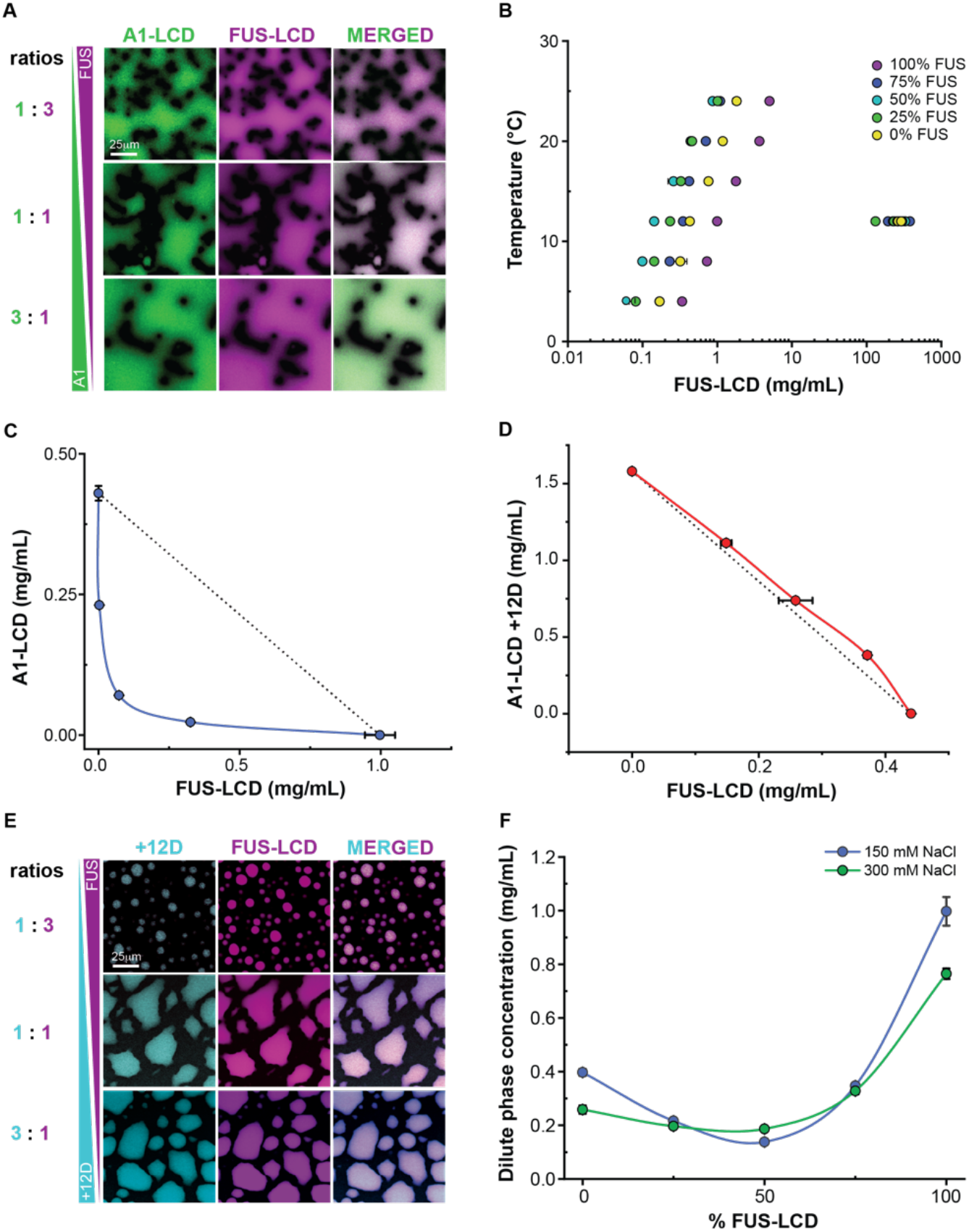
Data from *in vitro* phase separation assays of mixtures of PLCDs agree with simulation predictions. **(A)** Confocal fluorescence microscopy images of mixtures of FUS-LCD and A1-LCD at various mass concentration ratios. **(B)** Measured binodals of mixtures of FUS-LCD and A1-LCD. Concentrations correspond to the sum of the FUS-LCD and A1-LCD volume fractions. Percentages in the legend indicate the mass concentration ratio of FUS-LCD to A1-LCD. **(C),(D)** Measured 2-component binodals of mixtures of FUS-LCD and **(C)** A1-LCD or **(D)** A1-LCD +12D. Black dashed lines connect the intrinsic *c*_sat_ of FUS-LCD to the intrinsic *c*_sat_ of A1-LCD or A1-LCD +12D and are shown to indicate the expected binodal shape if heterotypic interactions are on par with homotypic interactions. **(E)** Confocal fluorescence microscopy images of mixtures of FUS-LCD and A1-LCD +12D at various mass concentration ratios. **(F)** Measured dilute phase total protein concentrations of mixtures of FUS-LCD and A1-LCD at various mass concentration ratios. Data are shown for mixtures in 150 mM NaCl or 300 mM NaCl. Where shown, error bars indicate standard deviations about the mean across at least three replicates.

As in the simulations, a 1:1 mixture of A1-LCD and FUS-LCD undergoes phase separation at a lower concentration than either protein on its own (**Fig. 3B**). Recasting the results as a twodimensional phase diagram shows the same pattern as the simulations, where the dilute arm of the phase boundary has a concave shape (**Fig. 3C**). These results suggest that complementary electrostatic interactions contribute to enhance co-phase separation of FUS-LCD and A1-LCD. In contrast, and in agreement with computational predictions, the dilute arm of the measured phase diagram for mixtures of the FUS-LCD and A1-LCD +12D system has a more convex shape (**Fig. 3D**).

Notably, although mixtures of FUS-LCD and A1-LCD +12D have a weakened driving force for phase separation relative to FUS-LCD and A1-LCD, the two proteins still co-localize into a single dense phase (**Fig. 3E**). We measured how the total dilute phase protein concentration varied as a function of salt concentration and the ratio of FUS-LCD-to-A1-LCD (**Fig. 3F**). Increasing the salt concentration from 150 mM NaCl to 300 mM NaCl resulted in decreased *c*_sat_ values for either protein on its own, but an increased *c*_dilute_ value for a 1:1 mixture. We rationalize this as follows: In solutions with only one type of PLCD, all proteins have the same sign and magnitude of charge. Accordingly, the intermolecular electrostatic interactions will be repulsive. Increasing the salt concentration screens repulsive interactions, thereby lowering the intrinsic *c*_sat_ values. In contrast, attractive electrostatic interactions in the 1:1 mixture will be weakened due to charge screening, and this leads to an increase in *c*_dilute_ for the mixture as salt concentration is increased. These results suggest an interplay between homo- and heterotypic aromatic interactions with intermolecular electrostatic interactions in mixtures of FUS- and A1-LCD.

### Tie lines and their slopes help uncover the relative contributions of homotypic versus heterotypic interactions to phase separation in mixtures of PLCDs

In a system that forms precisely two coexisting phases, the generalized tie simplex (39) for a given composition will be a tie line. The tie line is a straight line that connects points of equal chemical potentials and osmotic pressures on the dilute and dense arms of phase boundaries (69). The signs and magnitudes of the slopes of tie lines provide quantitative insights regarding the interplay between homotypic versus heterotypic interactions.

In a system with macromolecules A and B that undergo co-phase separation from a solvent, we can map the full phase diagram on a plane with the concentration of A being the variable along the abscissa, and the concentration of B being the variable along the ordinate (**Fig. 4**). In these titrations, we fix the solution conditions including the temperature. The slope of the tie line will be unity if heterotypic interactions are the dominant drivers of phase separation or if the homotypic interactions are equivalent to each other while also being on par with the heterotypic interactions. In the A-B mixture, a tie line with a slope that is less than unity will imply that homotypic interactions of component A are the main drivers of phase separation. Conversely, if tie lines have slopes that are greater than unity, then the homotypic interactions among B-molecules are the stronger drivers of phase separation.

**Fig. 4:**
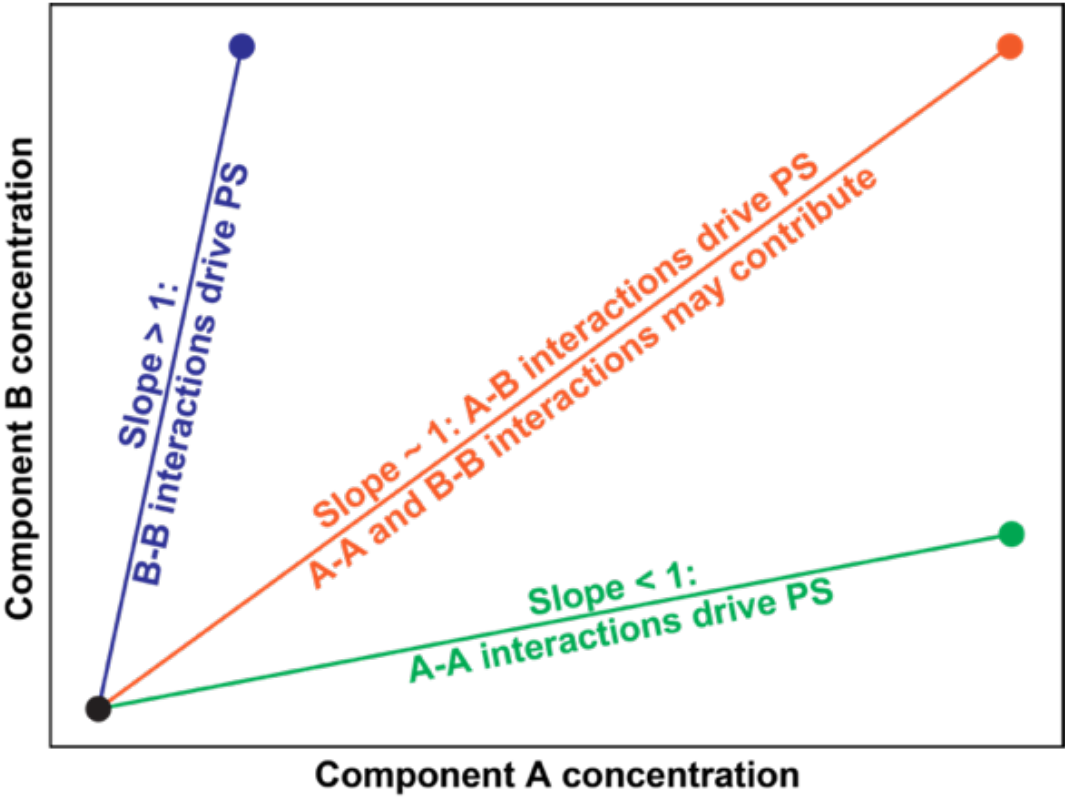
Schematic depicting how to interpret tie lines for a system of two components that undergo co-phase separation from the solvent to form precisely two coexisting phases.

If the total concentration of the mixture is *c*_tot_, we can extract tie lines by joining three points corresponding to *c*_dilute_, *c*_tot_, and *c*_dense_. Here, *c*_dilute_ and *c*_dense_ are the macromolecular concentrations in the coexisting dilute and dense phases, respectively. A single line connects the three points if and only if phase separation gives rise to precisely two coexisting phases. To test that this is the case, we plotted two sets of lines for each mixture *viz.*, one that joins *c*_dilute_ and *c*_tot_ and another that joins *c*_tot_ and *c*_dense_. If the slopes of the two lines are identical, then the system forms precisely two coexisting phases upon phase separation. The tie lines computed in this manner are shown in **Fig. 5A** and **5B** for simulations of different ratios of A1-LCD-to-FUS-LCD. The slopes of the two sets of tie lines are essentially equivalent to one another (**Fig. 5C**). We repeated this analysis for measured phase diagrams of the corresponding system *in vitro* and find similar values for the slopes of the tie lines (**Fig. 5D-F**).

**Fig. 5:**
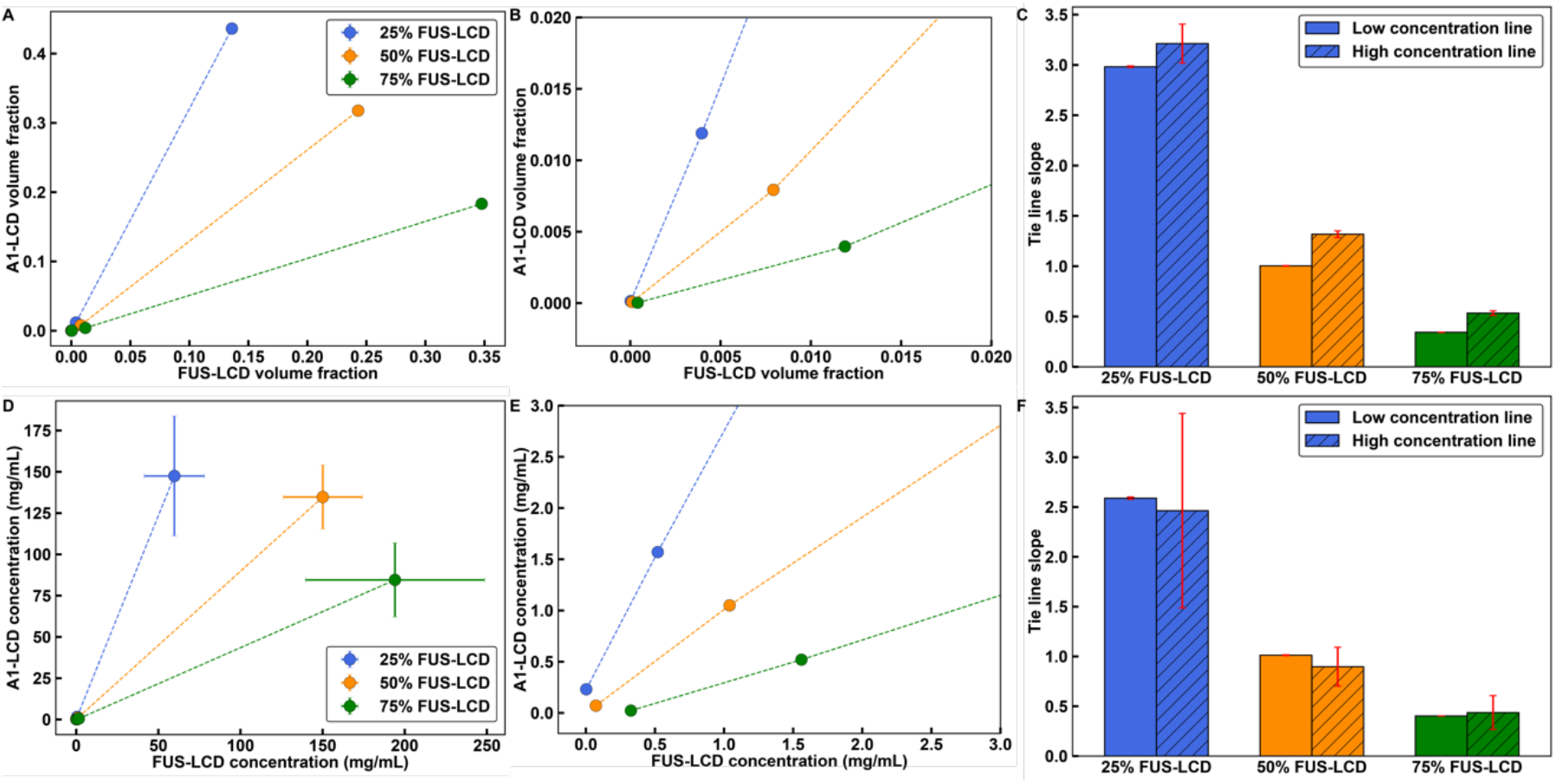
Tie lines of two-dimensional phase diagrams provide information regarding the interplay between heterotypic and homotypic interactions. Here, results shown in panels **(A)-(C)** are from simulations and those in panels **(D)-(F)** are from measurements. **(A), (B)** Tie lines extracted from simulations of phase separation in mixtures of FUS-LCD and A1-LCD. The simulation temperature is 49 in reduced units (see *SI Appendix*). **(A)** This plot includes the dilute phase concentrations, the total concentrations, and the dense phase concentrations. **(B)** This plots shows only the dilute phase concentration and total concentrations. **(C)** Computed slopes of the tie lines in **(B)** connecting dilute phase concentrations to the total concentrations and the total concentrations to the dense phase concentrations. The values are very similar to one another. **(D), (E)** Tie lines extracted from *in vitro* measurements of the FUS-LCD and A1-LCD system obtained at a temperature of 12°C. **(D)** Plot shows the dilute phase concentrations, the total concentrations, and the dense phase concentrations, whereas **(E)** focuses on the dilute phase concentration and total concentrations. **(F)** The slopes of the tie lines in **(E)** connecting dilute phase concentrations to the total concentrations and the total concentrations to the dense phase concentrations. Error bars in all panels are standard deviations about the mean across five replicates for simulations and at least three replicates for *in vitro* measurements. Total system concentrations do not have error bars. Otherwise, where error bars are invisible, they are smaller than the marker size.

In computations and measurements, the 1:1 mixtures of A1-LCD and FUS-LCD have tie lines with slopes that are essentially one. Therefore, phase separation involves changes in concentrations from the dilute to the dense phase that are similar for both sets of molecules. This suggests that heterotypic interactions are the dominant drivers of phase separation in 1:1 mixtures. Conversely, in a mixture with a 3:1 ratio of A1-LCD-to-FUS-LCD, the slope of the tie line is significantly greater than one. The implication is that homotypic interactions among A1-LCD molecules play a dominant role in driving phase separation along this tie line. Likewise, in a mixture with a 1:3 ratio of A1-LCD-to-FUS-LCD molecules, the slope of the tie line is significantly less than one, highlighting the importance of homotypic interactions among FUS-LCD molecules along this tie line.

Next, we asked if the tie lines for the mixture of FUS-LCD and A1-LCD +12D would differ from those of FUS-LCD and A1-LCD. FUS-LCD has a lower intrinsic *c*_sat_ than A1-LCD +12D. First, we plotted the low concentration arms obtained from *in vitro* measurements (**Fig. 6A**). Following Qian et al. (69), the agreement between the two sets of tie lines in **Fig. 5F** suggests that we can infer the slope of the whole tie line based on the slope of the line that connects *c*_dilute_ to *c*_tot_. Unlike the mixture of A1-LCD and FUS-LCD, we find that for the FUS-LCD and A1-LCD +12D mixture, the slope of the tie line for the 1:1 mixture is significantly less than one (**Fig. 6B**). This suggests that along this tie line, the homotypic interactions among FUS-LCD interactions are major drivers of phase separation. This effect is further enhanced for the mixture with a 3:1 ratio of FUS-LCD-to-A1-LCD +12D. Finally, for a mixture with a 1:3 ratio of FUS-LCD-to-A1-LCD +12D, the slope of the tie line is closer to unity, indicating that along this tie line, the interplay among heterotypic interactions, homotypic FUS-LCD interactions, and homotypic A1-LCD +12D interactions, are relatively well-balanced (**Fig. 6B**). Taken together, we find that when there are repulsive interactions between molecules in the mixture, the co-phase separation is driven by molecules that have stronger intrinsic driving forces for phase separation.

**Fig. 6:**
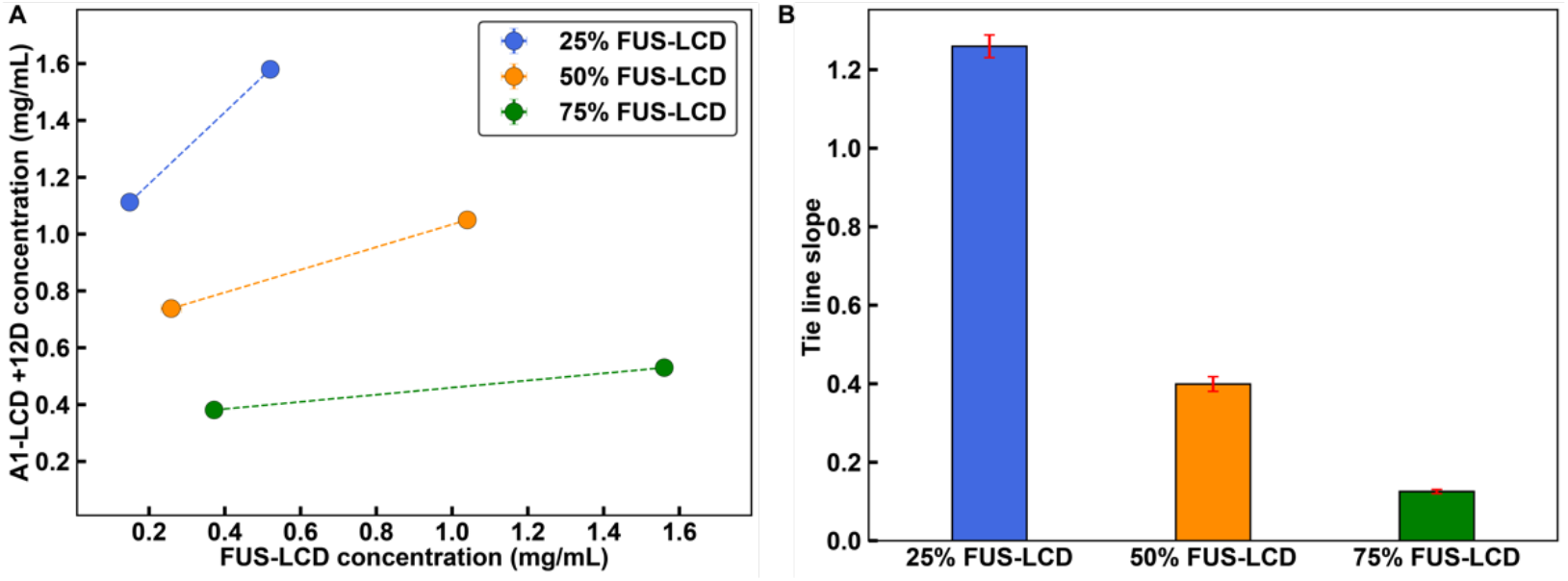
Weakening heterotypic interactions causes tie lines to be more sensitive to homotypic interactions. **(A)** Tie lines of the *in vitro* FUS-LCD and A1-LCD +12D system at a temperature of 4°C. These tie lines connect dilute phase concentrations to total concentrations. **(B)** Tie line slopes from **(A)**. Error bars in all panels are standard deviations about the mean across at least 3 replicates. Total system concentrations do not have associated error bars. Otherwise, where error bars are invisible, they are smaller than the marker size.

### How are condensate interfaces influenced by the balance of homotypic versus heterotypic interactions?

Next, we followed recent approaches (55) and computed radial density distributions, and logistic fits to these distributions (70) for a 1:1 mixture of A1-LCD and FUS-LCD (**Fig. 7A**). We find that the concentration of A1-LCD is greater than that of FUS-LCD in the dense phase and less than that of FUS-LCD in the dilute phase. This is because the intrinsic *c*_sat_ of A1-LCD is less than that of FUS-LCD. The thickness of the interfacial region predicted by the logistic fit to the radial density distribution for FUS-LCD is larger than that of A1-LCD. This is because FUS-LCD is ~1.6 times longer than A1-LCD.

**Fig. 7:**
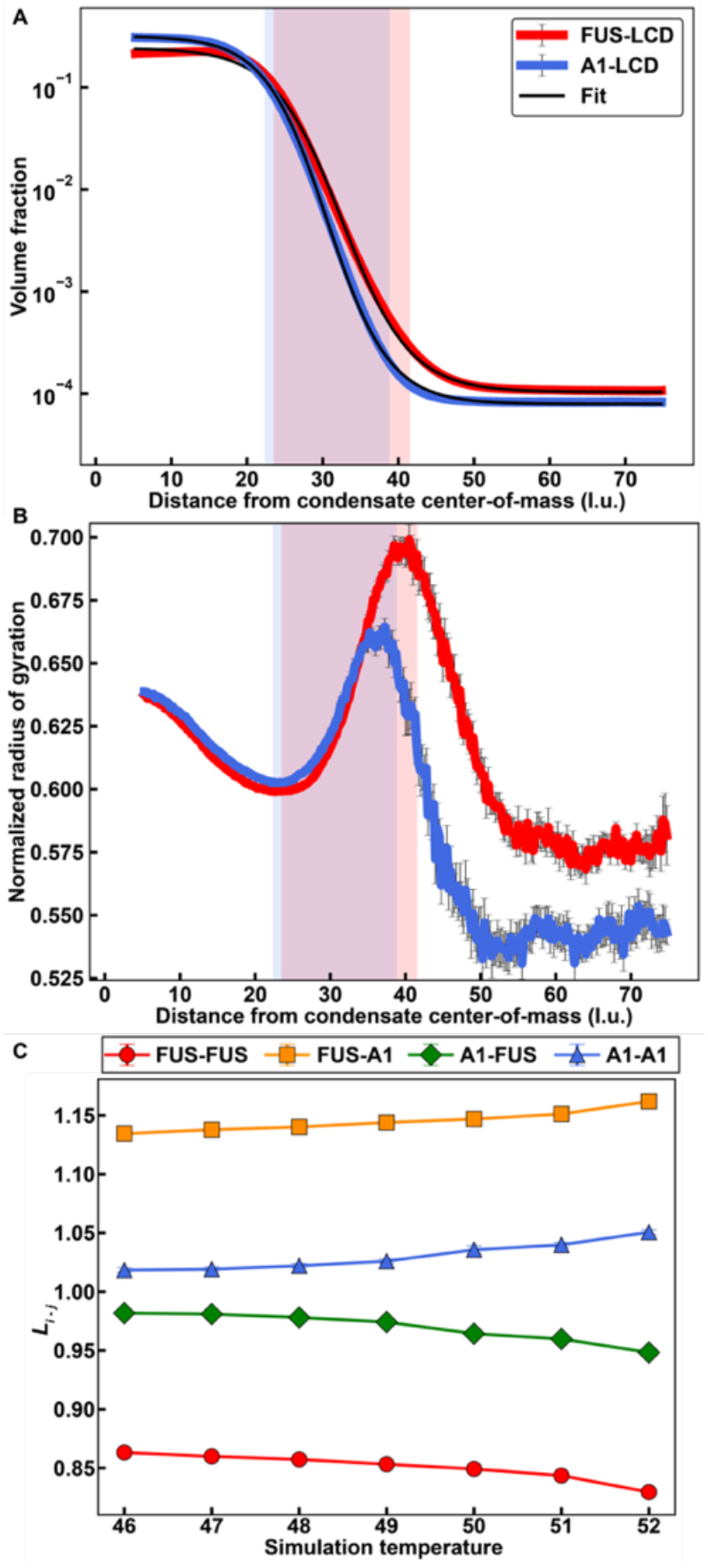
Simulated mixtures of PLCDs display complex interfacial features and internal organizations. **(A)** Radial density plots of simulated FUS-LCD and A1-LCD in a 1:1 mixture. Solid black curves indicate logistic fits to each radial density profiles. **(B)** Radius of gyration normalized by √*N*, where *N* is the chain length, as a function of the distance from the condensate center-of-mass. **(C)** Values of *L_i–j_* (see text) as a function of simulation temperature for a 1:1 mass concentration mixture of FUS-LCD and A1-LCD. In **(A)** and **(B)**, simulations were performed at a simulation temperature of 49, and translucent rectangles indicate the interfacial regions predicted by the corresponding logistic fit of FUS-LCD (red) and A1-LCD (blue). Error bars in all panels are standard errors about the mean across five replicates.

We analyzed the ensemble averaged radius of gyration, *R*_g_, of each species, normalized by √*N*, where *N* is the protein length (**Fig. 7B**). In the dense phase, each species has a similar value of *R*_g_ / √*N*, because the dense phase is an equivalently better solvent for both species when compared to the dilute phase, where the solvent is relatively poor for both species (55). In the dilute phase, *R*_g_/√*N* is slightly larger for FUS-LCD than for A1-LCD, and this is in accordance with the weaker homotypic interactions among FUS-LCD molecules. At the interface, both species show the predicted chain expansion (55).

Next, we probed the internal organization of A1-LCD and FUS-LCD molecules with respect to one another. To quantify this, we deployed a crosslinking parameter, *L_i–j_*, to determine the relative likelihood that protein species *i* interacts with species *j*, given a fixed total number of proteins from each species in the condensate. Details of how *L_i–j_* are computed are described in the *SI Appendix.*

Values of *L_i–j_* that are close to one indicate that species *i* interacts with species *j* in a manner that is proportional to the number of *i* and *j* proteins in the condensate i.e., the proteins are randomly mixed. In contrast, values of *L_i–j_* that are greater than or less than one point to a nonrandom organization of different molecules within the condensate. If *L_i–j_* > 1, then species *i* is more likely to interact with species *j* than would be expected from random mixing. Conversely, if of *L_i–j_* < 1, then species *i* is less likely to interact with species *j* than would be expected for a random mixture.

We calculated *L_i–j_* for simulations of FUS-LCD and A1-LCD at 1:1 ratios and a series of temperatures below the critical phase separation temperature and found the following trends: In general, FUS-LCD molecules are more likely to interact with A1-LCD molecules than with other FUS-LCD molecules. In contrast, A1-LCD molecules show relatively equal preferences for interacting with FUS-LCD or A1-LCD molecules. As the temperature increases, both FUS-LCD and A1-LCD molecules show an increased preference for interacting with A1-LCD molecules. This is because the critical temperature for FUS-LCD is lower than the critical temperature of A1-LCD (55). The results outlined in **Fig. 7** set up predictions for internal organization and interfacial properties based on protein length and the interplay between homotypic and heterotypic interactions.

### Uncovering rules for internal and interfacial organization in mixtures that form co-localized condensates

To uncover general rules for how the interplay of homotypic versus heterotypic interactions influences the internal organizations of condensates, we chose simpler systems and investigated these in a series of separate simulations. These homopolymeric systems have been shown to be reasonable facsimiles of PLCDs (55). We performed simulations of 1:1 mixtures of two homopolymers, designated as A and B. Each polymer has 150 beads, and we varied the strengths of homotypic (A-A and B-B interactions) and heterotypic (A-B interaction) energies. For each model, we calculated *L_i–j_* at various temperatures corresponding to the two-phase regime of the composite system.

First, we set all homotypic and heterotypic interactions to be equivalent. In this scenario, we find that *L_i–j_* equals one for all species combinations and at all temperatures, suggesting A and B are randomly mixed within the condensate (**Fig. 8A**). However, if heterotypic interactions are stronger than homotypic interactions, then we observe an increase in A-B contacts, indicating that chains within the condensate are organized to maximize heterotypic interactions (**Fig. 8B**). Setting the homotypic interactions to be stronger than heterotypic interactions give rise to internally demixed condensates with an A-rich region forming an interface with a B-rich region (**Fig. 8C**). Similar findings were recently reported by Welles et al., (71). Finally, adding asymmetry to the energetics by keeping A-B and B-B interactions equivalent, but weakening A-A interactions leads to a strong preference for A molecules to interact with B molecules over other A molecules, whereas B molecules show indifference at low temperatures, but a preference for other B molecules at high temperatures (**Fig. 8D**). These observations are akin to those made for mixtures of A1-LCD and FUS-LCD molecules (**Fig. 7C**).

**Fig. 8:**
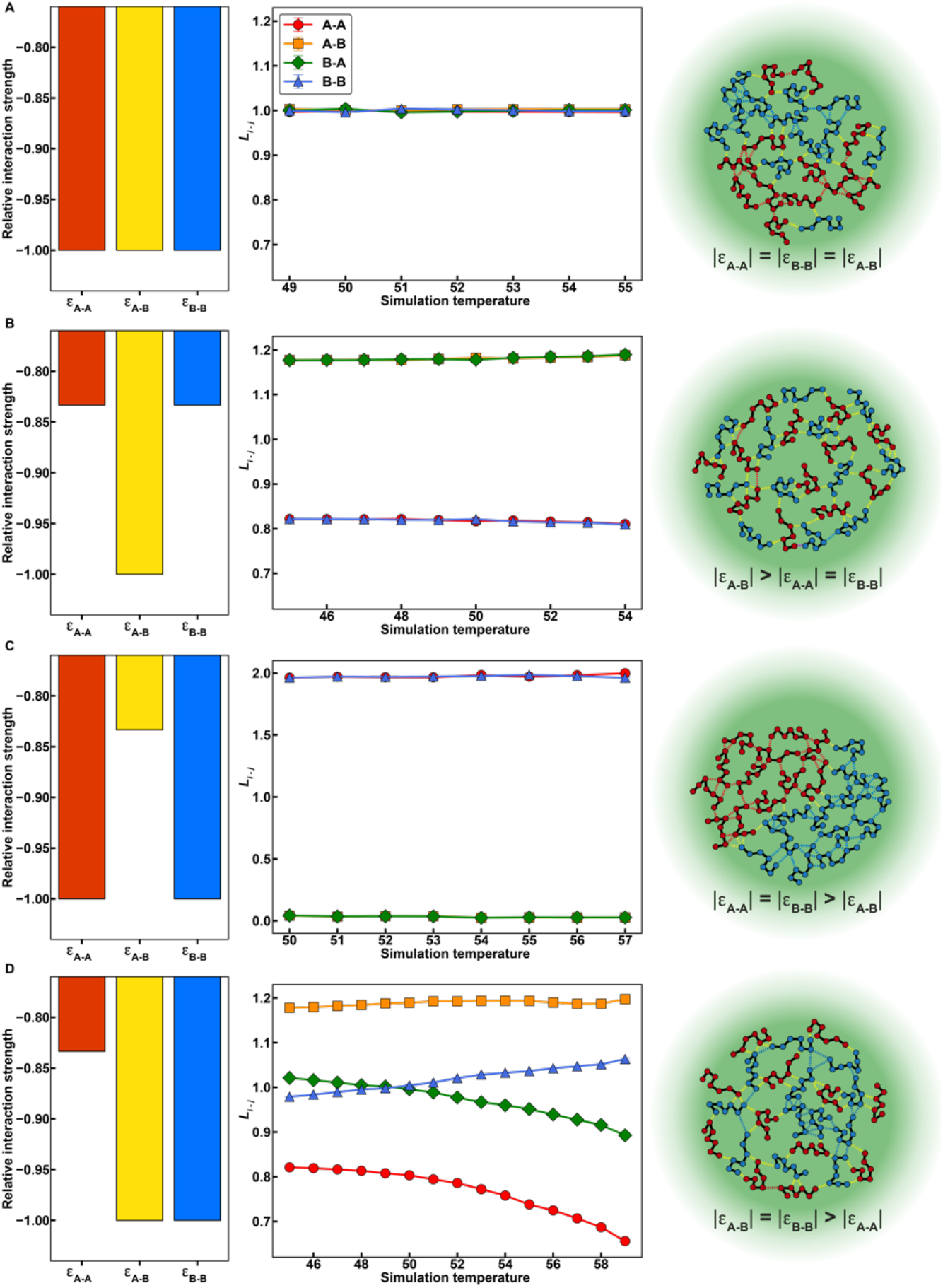
Internal condensate organizations of polymer mixtures depend on the interplay of homotypic vs. heterotypic interactions. **(A-D)** Relative interaction energies (left column), values of *L* (see text) as a function of simulation temperature (central column), and a schematic depicting the predicted condensate organization (right column) for 1:1 mixtures of polymers A and B with various interaction matrices. **(A)** Homotypic and heterotypic interactions are all equivalent. **(B)** Heterotypic interactions are stronger than homotypic interactions. **(C)** Homotypic interactions are stronger than heterotypic interactions. **(D)** Homotypic A-A interactions are weaker than homotypic B-B interactions and heterotypic A-B interactions, which are equivalent. Error bars for the values of *L* are standard errors about the mean across 5 replicates, though they are typically smaller than the marker size.

The lengths of macromolecules will influence the material properties (72) and interfacial features of condensates (55). Therefore, we asked how sequence length affects internal and interfacial features of condensates using simulations of homopolymers of various lengths. We performed simulations with equivalent mass concentrations for mixtures of homopolymers of lengths 150 (H150) and 300 (H300) where all homotypic and heterotypic interactions were set to be equal (**Fig. 9A**). Calculated phase diagrams show that H300 has a lower *c*_sat_ than H150 (*SI Appendix*, **Fig. S2A**). In addition, analysis of *L_i–j_* for the mixture shows that there is a strong preference for interacting with H300 above *T* ~ 56 in reduced units. Beyond this temperature, H150 no longer phase separates on its own (**Fig. S2A**). Radial density distributions show that H300 has a stronger preference for the dense phase and a weaker preference for the dilute phase when compared to H150. Logistic fits of the radial densities, used to determine interfacial regions corresponding to each homopolymer, show that the interfacial region is wider for H300 than for H150. We also analyzed the normalized radius of gyration and found that H300 has a greater degree of chain expansion in the interface when compared to H150. This is true even though it is more compact than H150 in the dense and dilute phases. Along the radial coordinate from the center of the condensate, the location within the interface that corresponds to the peak of chain expansion of H300 is shifted closer to the dilute phase when compared to the corresponding peak for H150.

**Fig. 9:**
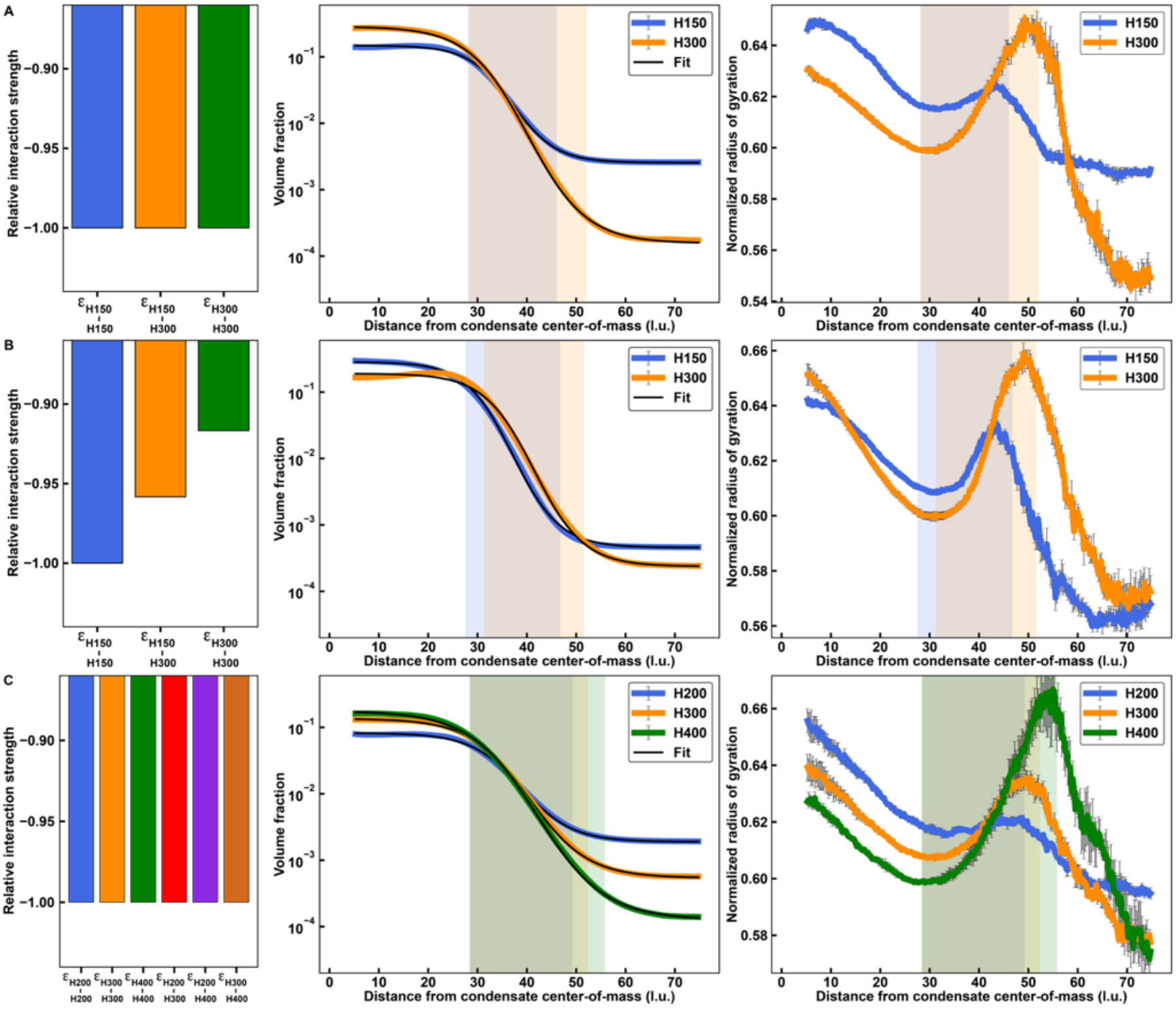
Interfacial features of multi-component condensates depend on polymer lengths and relative interaction energies. **(A-C)** Relative interaction energies (left column), radial density plots and associated logistic fits (central column), and normalized radius of gyration plotted against the distance from the condensate center-of-mass (right column) for equal volume fraction mixtures of two or three homopolymers of various lengths and interaction matrices. **(A)** Mixture of a 150-monomer polymer and a 300-monomer polymer where homotypic and heterotypic interactions are equivalent. **(B)** Mixture of a 150-monomer polymer and a 300-monomer polymer where relative interaction strengths are chosen such that the phase diagrams of the two polymers overlay. **(C)** Mixture of a 200-monomer polymer, a 300-monomer polymer, and a 400-monomer polymer where homotypic and heterotypic interactions are equivalent. Error bars are standard errors about the mean across five replicates.

To assess the extent to which our findings reflect the stronger intrinsic driving forces for phase separation of longer homopolymers, we titrated the strengths of homotypic and heterotypic interactions of H300 and H150 so that the *c*_sat_ values of H300 and H150 were equivalent (**Fig. 9B** and *SI Appendix*, **Fig. S2B**). In mixtures of equivalent mass concentrations, the mixture of H150 and H300 molecules forms apparent core-shell structures that allow both H150 and H300 to maximize their interactions with H150 molecules, as shown by an analysis of *L_i–j_* (*SI Appendix*, **Fig. S2B**). In turn, the width of the interfacial region defined by H300 shrinks when compared to the results shown in **Fig. 9A**. The scaled *R*_g_ values for H150 and H300 are similar in the dilute and dense phases. However, the locations and heights of the peaks of chain expansion show the same pattern as in **Fig. 9A**.

Finally, we asked if interfacial features vary when we extend to three-component systems. In simulations that use equal mass concentration of homopolymers of lengths 200, 300, and 400, with all interaction strengths being equal, we find that interfacial organizational features follow rules that were uncovered for binary mixtures. This is evident in comparisons of results shown in **Fig. 9C** and *SI Appendix*, **Fig. S2C** to those shown in **Fig. 9A**. Longer polymers make up more of the outer regions of the interface. They also show a greater level of expansion at the interface, and the peak of chain expansion shifts toward the dilute phase relative to that of the shorter polymers.

## Discussion

Condensates are characterized by distinctive macromolecular compositions (73–77). It is thought that macromolecular compositions, delineated into categories of molecules known as scaffolds, clients, regulators, and ligands, contribute to the functions of condensates (78–83). Here, we explored how the interplay between homotypic versus heterotypic interactions of PLCDs with at least cursorily similar compositional biases can influence the driving forces for phase separation. Our results set the stage for understanding how dynamical control over compositions in complex mixtures might be achieved and how this can influence condensate formation of bodies such as 40S hnRNP particles (84) or stress granules. There is also growing interest in understanding how the interplay of homotypic and heterotypic interactions impact the dynamically controlled (85) or purely thermodynamically influenced miscibility of condensates formed in multicomponent systems (71, 86–88).

Key findings that emerge from our investigations are as follows: In 1:1 binary mixtures of PLCDs, heterotypic interactions can enhance the driving forces for phase separation. This enhancement comes from complementary electrostatic interactions, thus suggesting a role for complex coacervation (43, 44) even in systems where fewer than 10% of the residues are charged. This highlights additional roles for spacer residues, which determine the solubility and the extent of coupling between associative and segregative transitions for individual PLCDs, while they determine the electrostatic complementarity of mixtures of PLCDs. Within condensates, the concentration of macromolecules will be above the overlap threshold (55). As a result, the interplay between homotypic and heterotypic interactions will contribute to the internal organization of molecules within condensates (89). Clearly, physicochemical control of cellular biomolecular condensates, which typically contain dozens of distinct macromolecular species, will involve a complex and dynamic interplay of different types of interactions between macromolecules. Our findings are reminiscent of the results of Espinosa et al., who used ultra-coarse-grained patchy colloid models for simulations of complex mixtures of associative macromolecules (90).

If phase separation gives rise to precisely two coexisting phases, then a 1-simplex (39) or tie line will connect the two points that define the coexisting phases. Each phase is defined by concentrations of macromolecules and solution components that enable equalization of osmotic pressures and chemical potentials i.e., partial molar free energies across the phases. The slopes of tie lines tell us about the relative contributions of homotypic and heterotypic interactions to phase separation for specific stoichiometric ratios of the macromolecules that drive phase behavior. Using a tie line analysis, we find that FUS-LCD and A1-LCD engage in strong heterotypic interactions when present in equal mass concentration ratios. However, when present in unequal ratios, homotypic interactions involving the predominant species become the primary driving force for phase separation. This behavior changes for FUS-LCD and A1-LCD +12D mixtures. For this system, phase separation in mixtures of equal mass concentration ratios is driven mainly by homotypic interactions among FUS-LCD molecules. Tilting the balance of interactions toward heterotypic interactions can be achieved by increasing the ratio of A1-LCD +12D to FUS-LCD. These results highlight the central importance of expression levels and stoichiometric ratios in mixtures of macromolecules (91).

We find that molecules within condensates are organized to maximize the most favorable interactions, be they homotypic or heterotypic. Accordingly, segregative transitions i.e., phase separation transitions drive the formation of condensates wherein associative interactions, i.e., physical crosslinking among favorably interacting molecules, are maximized (39). Further, molecules that engage in stronger interactions are more likely to be in the core of the condensate as opposed to in the interface. These findings highlight the crucial importance of the coupling of segregative and associative transitions as joint drivers of condensate formation and determinants of condensate internal structures.

In agreement with prior work (55), we find that polymers at the condensate interface are likely to adopt highly expanded conformations. We probed the effects of polymer lengths and found that the degree of expansion increases with polymer length. The strengths of homotypic versus heterotypic interactions and polymer length contribute to preferential localization of molecules at the interface, the degree of chain expansion at the interface, and the thickness of the interface. All polymers show an increase in their *R*g values near the condensate interface, and the precise location of the maximal expansion increase is controlled by the species-specific interface. In general, interfaces defined by longer polymers are thicker than those formed by shorter polymers. Interfacial thickness can be altered by modulating the strengths of homotypic and heterotypic interactions, allowing for finer control over spatial localization of each polymer species. The physical properties of interfaces (92) are likely to contribute to capillary forces of condensates (93), interactions between condensates and emulsifiers *in vivo* (94) or *in vitro* (95), and chemical reactions that are likely controlled at interfaces (96, 97).

Overall, our findings highlight the complex and designable properties (98–100) of biomolecular condensates formed by mixtures of PLCDs. These findings set the foundations for dissecting the phase behaviors and properties of condensates formed by *n*-nary mixtures of multivalent macromolecules. Our work highlights the promise of using simulations for modeling and designing phase behaviors in multicomponent systems (60, 87, 100–102).

## Materials and Methods

### Identifying condensate-associated prion-like low-complexity domains (PLCDs)

To identify the number of proteins with PLCDs that have been located within different cellular condensates, we used DrLLPS (103), a database of condensate-associated proteins. We culled all proteins associated with condensates with at least 40 known components. We then used the algorithm PLAAC (104) to identify proteins in the curated database that contain a PLCD. Within PLAAC, we specified a 50/50 weighting of background probabilities between *H. sapiens* and *S. cerevisiae.* This resulted in 89 distinct condensate-associated proteins with PLCDs. The sequences of these PLCDs were used for analyses that led to the results in **Fig. 1A-C**.

### Monte Carlo simulations using LaSSI

The coarse-grained model uses one lattice bead per amino acid residue and treats vacant sites as components of the solvent. In this way, solvent is afforded space in the system, but we do not parameterize any explicit interactions between solvent occupied sites and any other sites. Monte Carlo moves are accepted or rejected based on the Metropolis-Hastings criterion such that the probability of accepting a move is the min[1,exp(-Δ*E*/*k_B_T*)] where Δ*E* is the change in total system energy of the attempted move and *k_B_T* is the thermal energy, or simulation temperature. The energetic model used is the same as that described by Farag et al., (55). Here, we perform simulations of multi-component systems. We modified the mean-field electrostatic model, which was originally based on a single-component system. Instead of using a single NCPR value to determine the effect of electrostatics on a pairwise bead-bead interaction, we use the average NCPR of the chains to which the beads belong. The full set of Monte Carlo moves used in each type of simulation performed in this work are shown in **Table S1** of the *SI Appendix*. All the move sets and frequencies of moves are as described previously (55).

We performed multi-chain LaSSI simulations at various temperatures. The simulation temperatures are referenced in terms of units where we set *k_B_* = 1. Simulations involving FUS-LCD and A1-LCD used a 120×120×120 cubic lattice with periodic boundary conditions. Those involving homopolymers used a 150×150×150 cubic lattice with periodic boundary conditions. To speed up condensate formation, the simulations were initialized in a smaller 35×35×35 cubic lattice, which allows for significantly faster equilibration processes. The number of chains in each simulation was chosen to keep the total volume fraction of beads as close as possible to 0.016. Details of the sequences used in this study are shown in *SI Appendix* **Table S2**.

A total of 3×10^10^ MC steps was deployed for multi-chain simulations at each of the simulation temperatures. Simulations typically equilibrated after about 2×10^9^ steps, as determined by a plateauing of the total system energy. To be conservative, all simulation results were analyzed after the halfway point of 1.5×10^10^ steps. Multi-chain simulations were performed with five replicates, with each simulation initiated by a distinct random seed.

### Details of constructs used for in vitro measurements

We used the WT low-complexity domain (LCD) (residues 186-320) of human hnRNPA1 (UniProt: P09651; Isoform A1-A), in which the M9 nuclear localization signal had been mutated, substituting the PY motif with GS (referred to as A1-LCD); we also used a variant in which we further substituted several glycine and serine residues with aspartate residues increasing the aspartate content by 12 residues (A1-LCD^+12D^); and third, we used the WT LCD (residues 1-214) of FUS (UniProt: P35637). The gene sequences were synthesized as previously described (54), including a gene sequence coding for an N-terminal TEV cleavage site followed by the protein coding sequence of interest. The protein sequences are shown in *SI Appendix* **Table S2**. The three proteins were expressed and purified as previously described (53, 54, 66), and the purified proteins were stored in 6 M GdmHCl (pH 5.5), 20 mM MES at 4°C until they were buffer exchanged into phase separation buffer.

All other details of how the simulations were analyzed, and how the experiments were performed and analyzed are furnished in the *SI Appendix.*

## Data Availability

All experimental data and the code used to analyze simulations are available at the following Pappu Lab GitHub repository: https://github.com/Pappulab/Data-and-Analysis-for-PLCD-Mixtures. Details of the sequences used for analysis that yielded results in Fig. 1 are available via Github. Protein expression constructs for *in vitro* measurements are available from Addgene.

## Acknowledgments

This work was supported by grants from the US National Institutes of Health (R01NS121114) and the St. Jude Research Collaborative on the Biology and Biophysics of RNP granules to RVP and TM, and the Air Force Office of Scientific Research (FA9550-20-1-0241) to RVP.

## Supporting Information Appendix

### Additional Materials and Methods

#### Simulation analyses

The main text provides a summary of how the LaSSI simulations were setup. We identified the presence of a phase boundary and estimated dilute phase concentrations, dense phase concentrations, interface midpoints, and interface widths using a logistic fit of the radial density (1). To calculate concentrations, we used the exact prior for a cubic lattice with a given size. To determine chain expansion through the interface, we used radial shells with thickness of 1/4 of a lattice unit. When calculating the radial bins for the chains, rather than using the center-of-mass of a chain and counting each chain one time, we independently counted each bead in the chain using the radial bin of the bead and the radius of gyration of the corresponding chain. This accounts for the fact that a single chain can span multiple bins. The contribution of each bin is weighted by the number of beads of a chain that belong to the bin.

#### Crosslinking analysis

In this work, we introduce a crosslinking parameter, *L_i–j_* that quantifies the relative likelihood that a molecule from species *i* interacts with a molecule from species *j.* To calculate *L_i–j_*, we first calculate *f_i–j_*, the total number of intermolecular contacts between species *i* and *j* within the condensate (*N_i–j_*), divided by the total number of intermolecular contacts between species *i* and all other species:

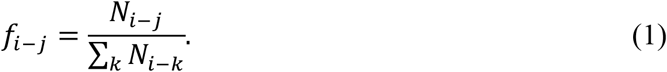

Next, we calculate *f*_*j*|*i*_, the total number of proteins of species *j* in the condensate from the perspective of a molecule of species *i* (*N*_*j*|*i*_), divided by the total number of molecules in the condensate from the perspective of a molecule of species *i*:

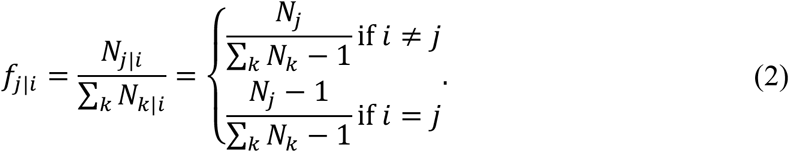

We subtract 1 from the calculation if *i* = *j* to account for the fact that we are only interested in intermolecular interactions. Accordingly, we ignore the protein from whose perspective we are calculating the given value. Finally, we take the ratio of these two values to calculate *L_i–j_*:

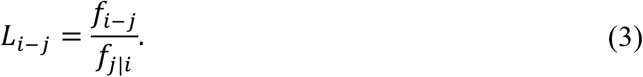

Values of *L_i–j_* close to 1 indicate that species *i* interacts with species *j* in a manner that is proportional to the number of *i* and *j* proteins in the condensate *i.e.*, the proteins are randomly mixed. In contrast, values greater than 1 indicate that species *i* is more likely to interact with species *j* than would be expected from random mixing, and values less than 1 indicate that species *i* is less likely to interact with species *j* than would be expected.

#### Phase separation assays in vitro

Samples were prepared as described in the Materials and Methods section of the main text. The buffer exchange was achieved in two-steps. The total protein concentration was kept at 2.1 mg/mL across all mixed conditions to be consistent with the ratios used in the simulations. As an example, for the 50% FUS ratio, enough A1-LCD was added to bring its final concentration to 1.05 mg/mL, and enough FUS-LCD was added to bring its final concentration to 1.05 mg/mL for a total protein concentration of 2.1 mg/mL. The protein solutions were mixed in 20 mM HEPES (pH 7.0), and 3 M NaCl in 20 mM HEPES (pH 7.0) was spiked into the solution to bring the final NaCl concentration to 150 mM. The samples were incubated at the desired temperatures for 20 min, then centrifuged at this temperature for 5 min at 12,000 rpm to separate the solution into dilute and dense phases. The dilute phase was then transferred to HPLC vials for analysis. A sample of the dense phase was taken for one temperature and diluted 1000-fold with 6 M GdmHCl (pH 5.5), 20 mM MES. The vials of the separated phases were then analyzed by analytical HPLC to assay the component protein concentrations in each phase.

#### Quantification of coexistence concentrations of mixtures of A1-LCD and FUS-LCD using analytical HPLC

We determined the saturation dilute and dense phase concentrations (*c*_sat_-*c*_dilute_ and *c*_dense_, respectively) for all A1-LCD and FUS-LCD mixtures using analytical HPLC (2). Before analyzing the mixtures on the HPLC we ensured that A1-LCD and FUS-LCD elute separately and with sufficient resolution for peak integration. To determine the corresponding saturation, dilute and dense phase concentrations of each component, we measured a standard curve for each component. The standard curve for each protein was fitted to equation 1 in Bremer et al., (2) which was then used to determine *c*_dilutesat_ and *c*_dense_ for each component. At least three replicates per condition were measured.

#### Fluorescent labeling of protein samples

To test for localization of protein species within condensates, eEach variant was fluorescently labeled at the N-terminus. WT A1-LCD was labeled with Alexa Flour488 (NHS Ester; ThermoFisher), A1-LCD +12D with LD-555 (NHS Ester; Lumidyne Technologies), and FUS-LCD was labeled with LD-655 (NHS Ester; Lumidyne Technologies). The protein samples were labeled under denaturing conditions, in 50 mM Phosphate buffer (pH 7.0), 6 M GdmHCl, and quenched with 20mM Tris.

#### Confocal fluorescence microscopy

The extent of co-localization of A1-LCD and FUS-LCD for the different A1-LCD and FUS-LCD mixtures was examined using a Zeiss LSM 980 Airyscan 2. Samples were prepared as described in the phase separation assay, but each protein was doped with a fluorescently labeled fraction of itself. 2 μL of the mixture of interest was placed between two coverslips sandwiched with 3M 300 LSE high-temperature double-sided tape (0.34 mm) with a window for microscopy cut out. All measurements were performed in 20 mM HEPES (pH 7.0), 150 mM NaCl at room temperature, and image analysis was performed using Fiji (Version 2.1.0/1.53o) (3).

**Supplementary Table S1:** Monte Carlo move sets for simulations.

| Move Sets | Probability of Choosing a Move for Multi-Chain Simulations |
| --- | --- |
| Local Moves | 0.51 |
| Multi-local Moves | 0.17 |
| Slithering Snake Moves | 0.03 |
| Translational Moves | 0.03 |
| Pivot Moves | 0.09 |
| Double Pivot Moves | 0.17 |

**Table S2:**
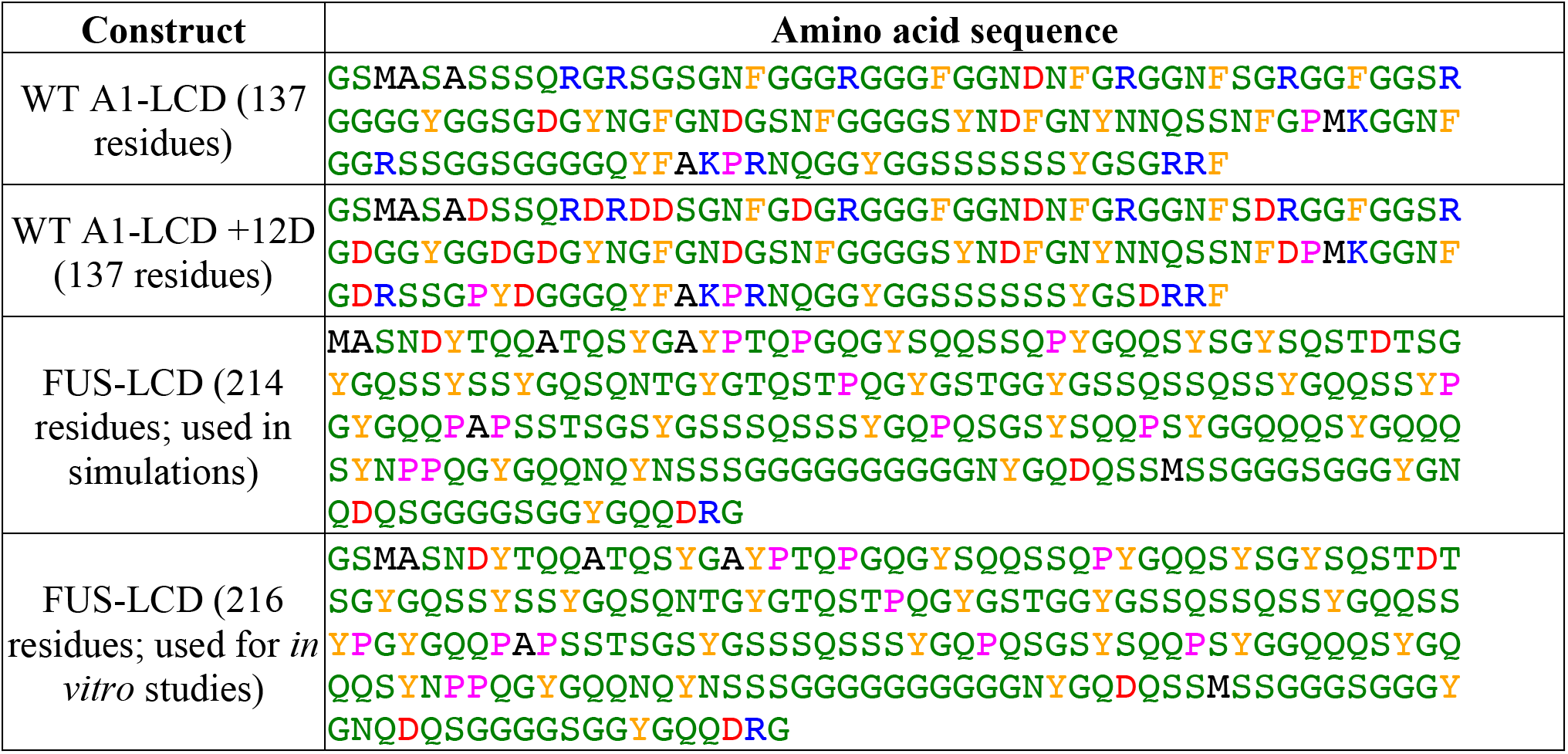
Protein constructs and their associated sequences. The FUS-LCD sequences used in simulations differs from that used in *in vitro* studies due to an N-terminal GS overhang that is a product of the *in vitro* purification method. The A1-LCD sequences used *in vitro* and in simulations contain this overhang, as these were the sequences used previously (1). Amino acids are color-coded as follows: Polar – green, positively charged – blue, negatively charged – red, aromatic – gold, proline – pink. Methionine and alanine are colored black.

**Fig. S1:**
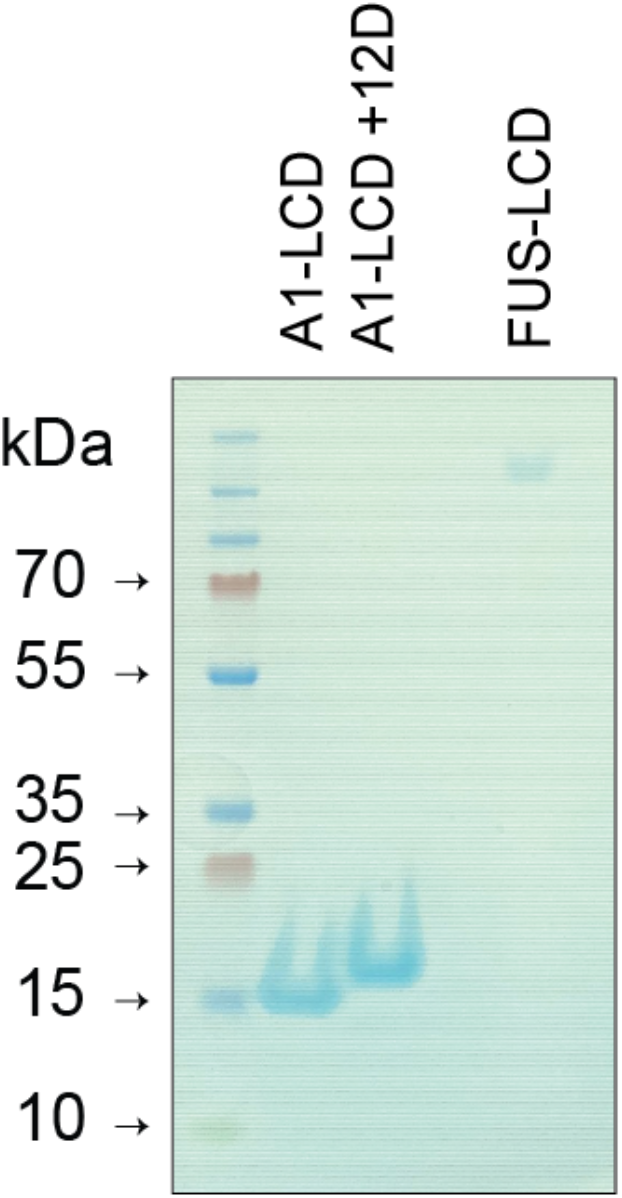
SDS-PAGE for A1-LCD, A1-LCD +12D, and FUS-LCD. FUS-LCD stains faintly with SimplyBlue stain due to low basic residue content. It also appears at a higher-than-expected molecular weight (MW). The intact mass analysis for FUS-LCD shows a MW of 21,697 Da, in line with the expected monomer mass. SEC and DLS data also confirm the expected size of the FUS-LCD.

**Fig. S2:**
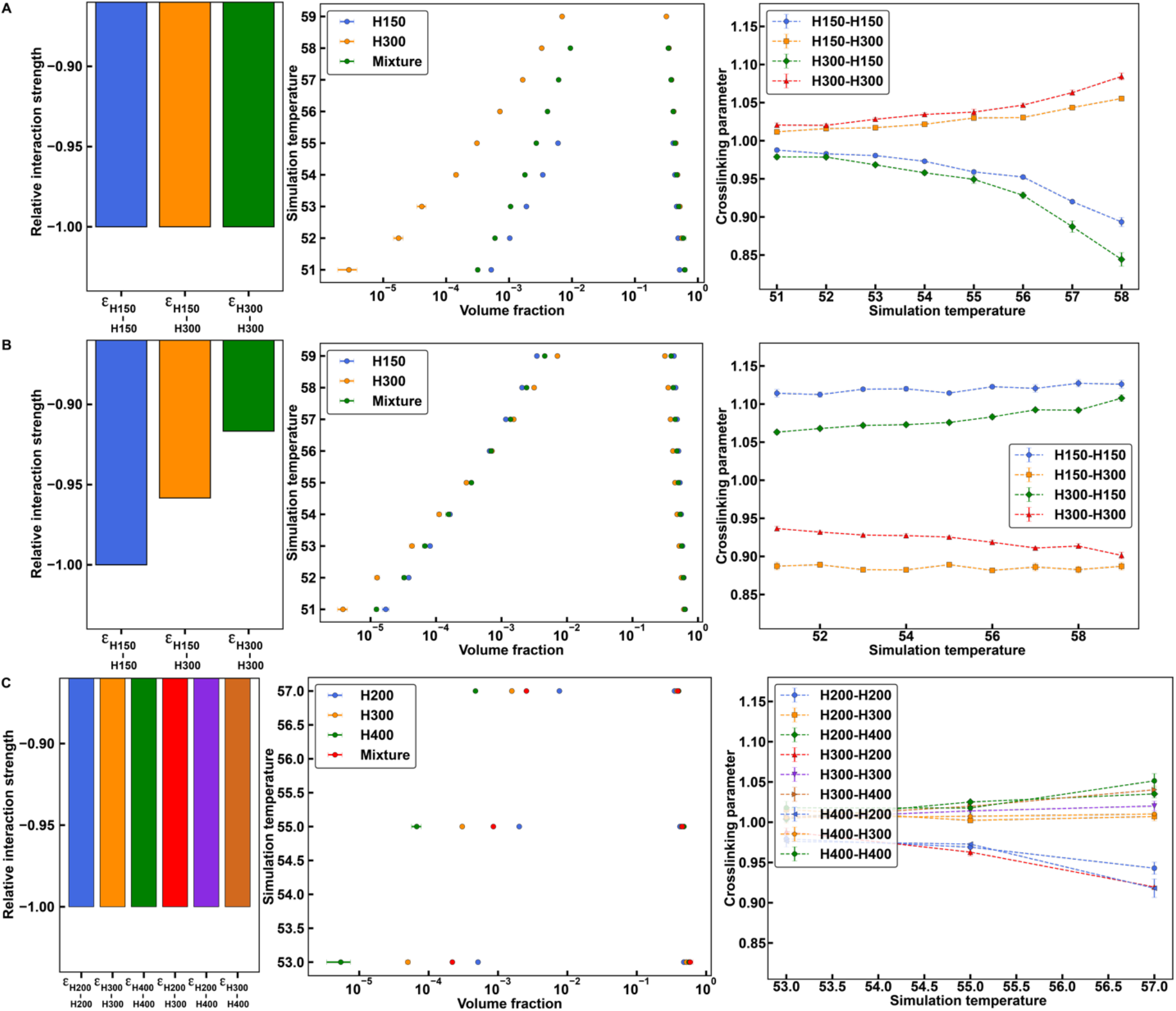
Phase diagrams and crosslinking parameters of mixtures of homopolymers with different lengths and interactions. **(A-C)** Relative interaction energies (left column), phase diagrams (middle column), and crosslinking parameters for equal volume fraction mixtures (right column) of 2 or 3 homopolymers of various lengths and interaction matrices. **(A)** Mixture of a 150-monomer polymer and a 300-monomer polymer where homotypic and heterotypic interactions are equivalent. **(B)** Mixture of a 150-monomer polymer and a 300-monomer polymer where relative interaction strengths are chosen such that the phase diagrams of the two polymers overlay. **(C)** Mixture of a 200-monomer polymer, a 300-monomer polymer, and a 400-monomer polymer where homotypic and heterotypic interactions are equivalent. Error bars are standard errors about the mean across five replicates.

## Notes

### Competing Interest Statement

The authors have declared no competing interest.

